# Evolutionary mysteries in meiosis

**DOI:** 10.1101/050831

**Authors:** Thomas Lenormand, Jan Engelstadter, Susan E. Johnston, Erik Wijnker, Christoph R. Haag

## Abstract

Meiosis is a key event of sexual life cycles in eukaryotes. Its mechanistic details have been uncovered in several model organisms, and most of its essential features have received various and often contradictory evolutionary interpretations. In this perspective, we present an overview of these often “weird” features. We discuss the origin of meiosis (origin of ploidy reduction and recombination, two-step meiosis), its secondary modifications (in polyploids or asexuals, inverted meiosis), its importance in punctuating life cycles (meiotic arrests, epigenetic resetting, meiotic asymmetry, meiotic fairness) and features associated with recombination (disjunction constraints, heterochiasmy, crossover interference and hotspots). We present the various evolutionary scenarios and selective pressures that have been proposed to account for these features, and we highlight that their evolutionary significance often remains largely mysterious. Resolving these mysteries will likely provide decisive steps towards understanding why sex and recombination are found in the majority of eukaryotes.

## Introduction

In eukaryotic sexual life cycles, haploid cells fuse to give rise to diploids, before diploid cells are converted back to haploids in a process known as meiosis. Meiosis reduces a cell's chromosome number by half, whilst also creating new allele combinations distributed across daughter cells through segregation and recombination. This genetic reshuffling reduces genetic associations within and between loci and is thought to be the basis of the success of sexual reproduction. Mechanistic studies of meiosis have been carried out in different fields, such as cell biology, genetics and epigenetics, encompassing a wide range of eukaryotes. However, these studies rarely focus on the evolutionary significance of meiotic mechanisms, rather mentioning them in passing and often in a simplified manner. In evolutionary biology studies, meiosis is often simplified and represented by random assortment of chromosomes and recombination maps expressing the probability of recombination events between ordered loci, with little attention to the molecular and cellular details. While these simplifications are legitimate and useful in many cases, the wealth of mechanistic findings being uncovered points to a considerable number of evolutionary puzzles surrounding meiosis that have yet to be resolved. Indeed, in the following perspective, we will show that close scrutiny of almost every aspect of meiosis will reveal “weird” features that constitute evolutionary mysteries.

## 1. The origins of meiosis

The origin of meiosis through gradual steps is among the most intriguing evolutionary enigmas [1,2]. Meiosis is one of the 'major innovations' of eukaryotes that evolved before their subsequent radiation over one billion years ago [3–5]. Extant eukaryotes share a set of genes specifically associated with meiosis, implying that it evolved only once before their last common ancestor [6,7]. Identifying the selective scenario that led to its early evolution is difficult, but clues can be obtained by determining (i) which mitotic cellular processes were re-used in meiosis (e.g. DNA repair through homologous recombination and possibly reduction), (ii) which selective steps were involved in the assembly of the full cellular process, and (iii) why different forms of meiosis were perhaps less successful.

### 1.1. The origin of ploidy reduction

A form of reductional cell division (a.k.a. ‘proto-meiosis') probably evolved in early asexual unicellular eukaryotes. Two scenarios for this have been proposed. The first is that diploidy accidentally occurred by replication of the nuclear genome without subsequent cell division (“endoreplication”) [8–12], and that returning to haploidy was selected for to correct this. Because either haploidy or higher ploidy levels may be favoured in different ecological situations [13,14], a variant of this scenario is that a proto-meiosis-endoreplication cycle evolved to switch between ploidy levels [5]. The resulting life cycle may have resembled modern ‘parasexual’ fungi in which diploid cells lose chromosomes in subsequent mitotic divisions, leading to haploidy via aneuploid intermediates [15]. Many other modern eukaryotes also increase and decrease their ploidy somatically, depending on growth stage or specific environmental stimuli [16]. The second scenario is that proto-meiosis evolved in response to the fusion of two haploid cells (“syngamy”), as in standard modern eukaryotic sexual life cycles. Syngamy may have been favoured because it allows recessive deleterious mutations to be masked in diploids [1,12]. A difficulty with this idea is that such masking may not be sufficient to favour diploidy in asexuals [17]. In a variant of this scenario, early syngamy evolved as a result of 'manipulation’ by selfish elements (plasmids, transposons) to promote their horizontal transmission [18]. In support of this view, mating-type switching (which can allow syngamy in haploid colonies) has evolved multiple times in yeasts and involves domesticated mobile genetic elements [19].

### 1.2 The origin of homologue pairing and meiotic recombination

Meiosis requires the correct segregation of homologues, which is achieved by homologue pairing at the beginning of prophase I (Fig. 1). This homology search is mediated by the active formation of numerous DNA double-strand breaks (DSBs) followed by chiasmata formation, but less well-known mechanisms of recombination-independent pairing also exist [20]. Non-homologous centromere coupling is also often observed at this stage, but the functional and evolutionary significance of this coupling is elusive [21]. In many species, chromosome pairing is further strengthened by 'synapsis', which is the formation of a protein structure known as the synaptonemal complex [22] and the pairing of homologous centromeres [21]. Chiasmata are then resolved as either crossovers (hereafter ‘COs’) resulting in the exchange of large chromatid segments, or noncrossovers (‘NCOs’), where both situations cause gene-conversion events [23]. The synaptonemal complex then disappears, and homologues remain tethered at CO positions and centromeres. The precise function of the synaptonemal complex is not entirely understood [20]; one possibility is that it may serve to stabilise homologues during CO maturation. Some pairing mechanism must be advantageous to ensure proper segregation of homologues, but the origins and selective advantage of extensive pairing, synapsis, gene conversion and recombination remain poorly understood [24].

**Figure 1.**
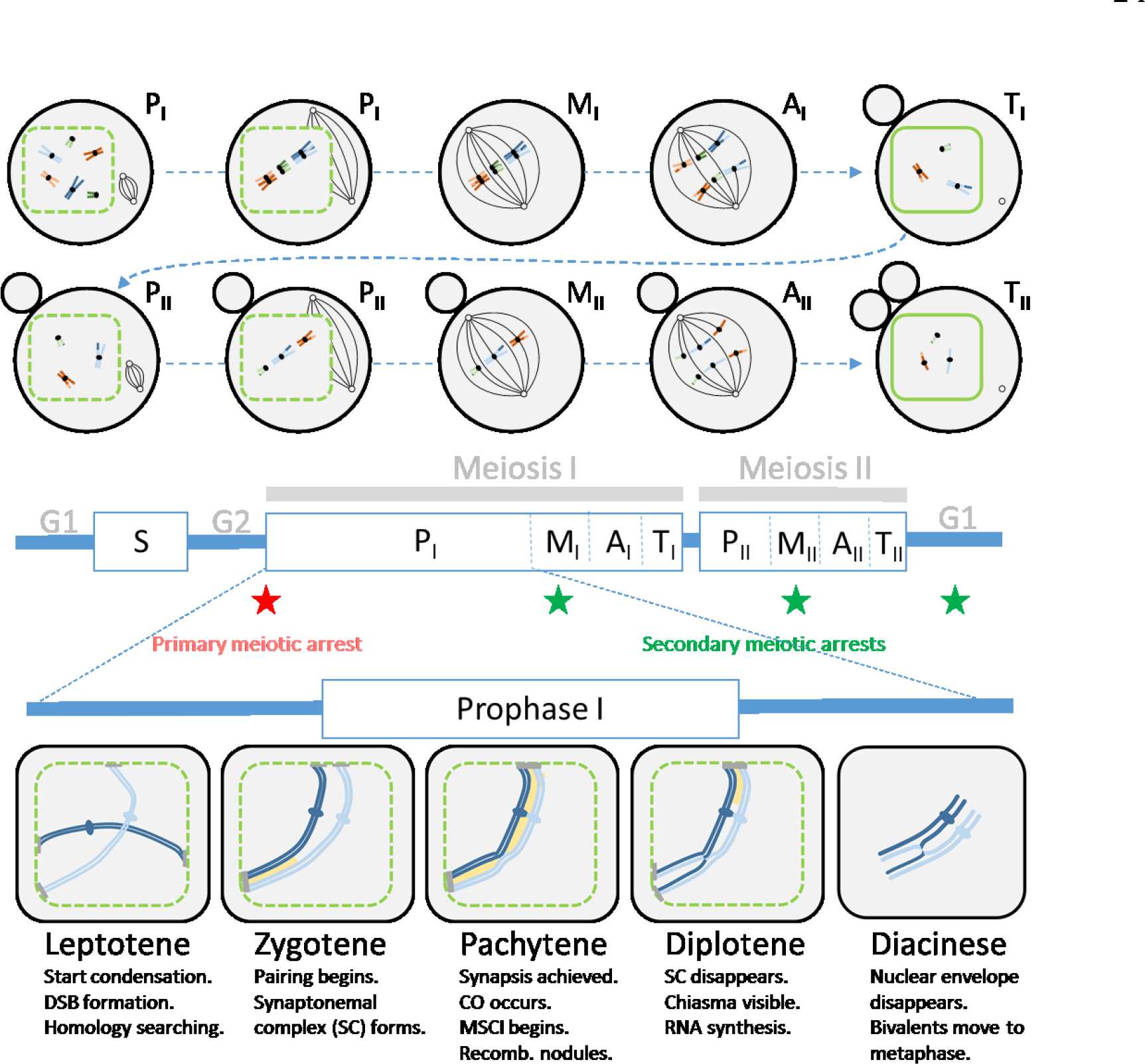
Schematic representation of the different steps in standard meiosis. The top panel illustrates the different phases of a typical female meiosis for each of the two meiotic divisions: prophase (P, with early and late prophase distinguished), metaphase (M), anaphase (A) and telophase (T). The nuclear membrane is indicated by the green contour (dashed when it starts fragmenting). The small black circles represent microtubule organizing centres and the black lines represent microtubules of the meiotic spindle. First and second polar bodies are shown as grey circles next to the oocyte (chromosomes inside the polar bodies are not shown). Homologous chromosomes are represented with the same colour with slightly different shades (e.g. orange and light orange). Homologues pair and segregate in meiosis I, then sister chromatids segregate in meiosis II. The middle panel shows the meiotic cell cycle. The timing of the primary meiotic arrest is indicated by a red star, while the timing of the most common secondary arrests in different organisms is indicated by green stars (see section 3.1). The lower panel indicates the important steps (DSB formation, crossing overs) occurring during prophase I. The synaptonemal complex is shown in yellow. Chromatin condenses in chromosomes throughout prophase I (only one pair of homologues is illustrated). In most999 species, telomeres attach to the nuclear envelope. The attachment plate is indicated by a1000 grey bar. MSCI refers to meiotic sex chromosome inactivation (see section 3.4).

Most evidence suggests that homologous recombination evolved long before meiosis, as it occurs in all domains of life and involves proteins that share strong homology [25,26]. One hypothesis is that meiotic pairing and extensive homologous recombination in meiosis evolved to avoid the burden and consequences of non-allelic ectopic recombination in the large genomes of early eukaryotes, which presumably had many repetitive sequences [9,27,28]. Such sequences may have been related to the spread of retrotransposons in early eukaryotes, of which many types are very ancient in eukaryotes, but absent in bacteria and archaea [29]. A second possibility is that recombination arose by the spread of self-promoting genetic elements exploiting the machinery of DNA repair and associated gene conversion [30]. Another hypothesis is that pairing and recombination initially arose as a way to repair mutational damage caused by increased oxidative stress due to rising atmospheric oxygen or endosymbiosis [7,31–33]. This scenario presupposes that DNA maintenance is inefficient in the absence of meiosis; however, prokaryotes (including archaea) have efficient repair mechanisms that involve recombination, but not meiosis [9]. In addition, this scenario does not fit well with the observation that a large number of DSBs are actively generated at the onset of meiosis [1,34].

### 1.3 The origin of two-step meiosis

A particular feature of meiosis is that it starts with chromosome doubling (S phase, see Fig. 1) before meiosis occurs (Fig. 2A). For ploidy reduction, the initial steps appear superfluous [35]. A simpler single-step cell division, without the initial DNA replication phase, could in principle achieve ploidy reduction (Fig. 2B). Recombination may not be a crucial difference between one-and two-step meiosis, as both can involve COs, even if with one CO, the two meiotic products carry recombinant chromosomes in one-step meiosis, whereas only two out of four are recombinant in two-step-meiosis [36]. Three hypotheses have been proposed to account for two-step meiosis. The first postulates that two-step meiosis better protects against particular selfish genetic elements (SGEs) that increase their transmission frequencies by sabotaging the meiotic products in which they do not end up (known as ‘sister killers’, distinct from the ‘sperm killers’ discussed below) [37]. In a two-step meiosis, there is uncertainty as to whether the reductional division is meiosis I or II, meaning that the sabotage mechanism has a much reduced efficacy. Microsporidia and red algae show specific modifications to meiosis that increase such uncertainty even more [38]. However, such sister killers are hypothetical, and theoretical studies based on assumptions about how different killers might act suggest that this mechanism does not inevitably promote the development of a two-step meiosis [39]. The second hypothesis is that sexual species with one-step-meiosis would be vulnerable to invasion by asexual mutants, and have thus gone disproportionally extinct in the past.

**Figure 2.**
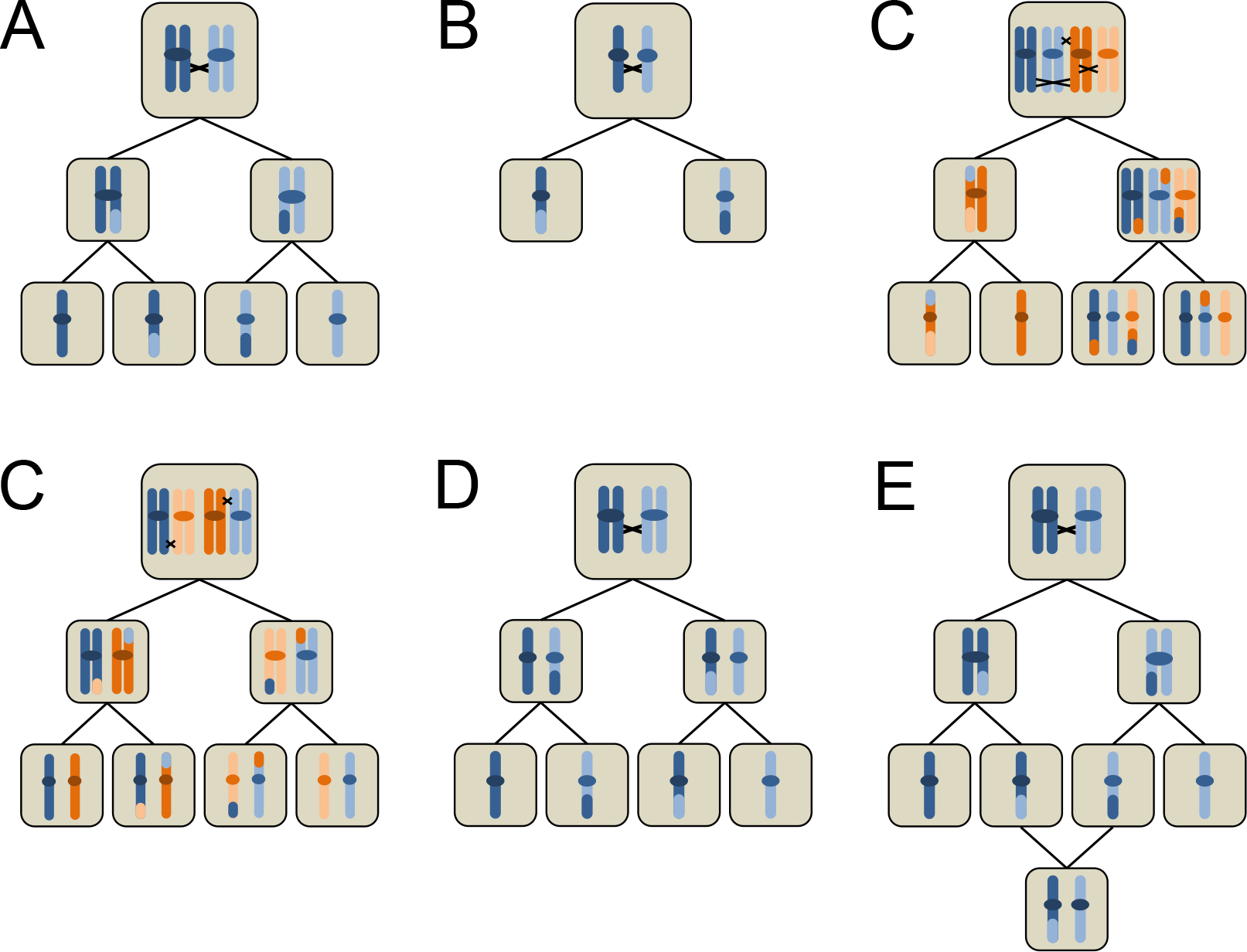
Schematic representation of meiosis and some of its modifications. (A) Regular meiosis. Following DNA replication, homologous chromosomes are separated in the first meiotic division, whereas sister chromatids are separated in the second division. COs result in chromosomes in the final meiotic products that carry genetic material from both homologous chromosomes. (B) Hypothetical “one-step” meiosis, in which DNA replication before entering meiosis is suppressed and therefore only a single meiotic division is required. (C) Multivalent formation in a neo-tetraploid. Blue and orange chromosome pairs are assumed to be identical or very similar so that pairing can occur. Chiasmata of one chromosome with three other chromosomes leads to mis-segregation. (D) Bivalent formation in a tetraploid with exactly one CO per chromosome. Chromosomes may pair randomly (leading to polysomic inheritance), but segregation proceeds normally. (D) Inverted meiosis, in which sister chromatids are separated in the first division and homologous chromosomes in the second division. Note that although centromeres are shown here for clarity, all described species consistently using inverted meiosis are holokinetic (no centromeres). (E) Central fusion automixis, a mechanism of producing diploid eggs that can then develop parthenogenetically without fertilisation. As a consequence of COs, heterozygosity may be lost with this mechanism in regions distal to the centromere.

Contrary to one-step meiosis, most automictic modifications of two-step meiosis involve a loss of heterozygosity with each generation (see section 2.3), which would cause expression of recessive and partially recessive deleterious mutational effects, and reduce the fitness of newly emerging asexual mutants [36]. Finally, a third hypothesis posits that a one-step-meiosis is more complex and thus less likely to evolve than a two-step-meiosis [9]. Mitotic and meiotic cell cycles start similarly with DNA replication in response to increasing cyclin-dependent kinase (CDK) activity. Two-step meiosis can be achieved simply by modulating CDK activity at the end of a cell cycle to add a second division event [40]. In contrast, a one-step meiosis would require extensive modification of the mitotic cycle. Despite earlier suggestions of its presence in some basal eukaryotes (protists) [8,41], there are presently no firm indications that one-step meioses exist in nature [38,42], although inverted meiosis (see below) is genetically similar to mitosis followed by single-step meiosis.

## 2. Secondary modifications of meiosis

Meiosis is remarkably conserved across eukaryotes. Nevertheless, in many species, variants exist that may offer insights into the evolutionary origins and mechanistic constraints of meiosis. Here, we discuss three of these modifications: meiosis in polyploids, inverted meiosis and meiosis in asexual organisms.

### 2.1 Meiosis and polyploidy

Polyploidy is surprisingly common in eukaryotes given the considerable problems it poses to meiosis [43–45]. In diploids, homologous chromosomes recognise each other and align to form bivalents during Prophase I, but when there are three or more chromosomes with sufficient homology, these chromosomes may all align to varying degrees, forming multivalents. This can occur when all chromosome sets originate from the same species (autopolyploidy), but also when polyploidy is a result of hybridisation (allopolyploidy). Multivalent formation is often associated with mis-segregation of chromosomes (Fig. 2C) as well as chromosomal rearrangements arising from recombination within multivalents, leading to reduced fertility and low-fitness offspring [e.g. 46,47,48]. These problems may be compounded in allopolyploids because recombination homogenises partially differentiated chromosomes, thereby further increasing the likelihood that they will pair [the ‘polyploid ratchet‘: 46].

Given these detrimental effects, the existence of successful polyploid species and lineages indicates that natural selection can often promote transitions from multi-to bivalents that will then segregate as in diploids (compare Figs. 2C & D) [e.g. 50,51]. However, how such transitions are achieved at the molecular level remains a mystery. Part of the answer seems to be a reduction in the number of COs, since multivalents can only form with at least two COs per chromosome [51–53]. This mechanism seems particularly important in autopolyploids and may be achieved through increasing CO interference (see section 4.2 for definition) [54]. Several candidate genes that may affect such modifications have been identified in the autotetraploid *Arabidopsis arenosa* [51,55]. In allopolyploids, there is evidence for genes that have been selected to strengthen the preferential pairing of homologous (i.e. of the same origin, rather than ‘homeologous’) chromosomes, including *phi* in hexaploid wheat [56]. This preferential pairing can also be achieved through reducing CO numbers, specifically those between homeologues; this could indirectly produce an increase in CO numbers and hence recombination rates between homologues [43]. Intriguingly, because most extant organisms have a history of polyploidy, many features of ‘standard’ meiosis such as CO interference may have been shaped by the problems involved in multivalent segregation.

Polyploidy with odd numbers of chromosome sets poses an even greater problem because aneuploid gametes are generally produced [e.g. 57]. However, there are some plant species where solutions to even this problem have evolved, and where odd-number polyploidy appears to persist in a stable manner. In these species, the problem of unequal segregation during meiosis is solved through exclusion of univalents in one sex but inclusion in the other, leading, for example, to haploid sperm and tetraploid eggs in pentaploid dog roses [58].

### 2.2 Inverted meiosis

In normal meiosis, homologous chromosomes are separated during meiotic division I, whereas sister chromatids are separated during meiosis II. Why meiosis generally follows this order is unknown, but interestingly, in some species meiosis takes place in the reverse order (Fig. 2E), including some flowering plants [59–61], mites [62], true bugs [63], and mealybugs [64]. All species with this ‘inverted’ meiosis described to date seem to have holocentric chromosomes (i.e. the kinetochores are assembled along the entire chromosome, rather than at localised centromeres). Inverted meiosis is viewed as a possible solution to specific problems of kinetochore geometry in such meiosis [65]. Yet, intriguing as they are, these systems provide little insight into why inverted meiosis is absent or very rare in monocentric species.

It is conceivable that a reverse order of divisions would make meiosis more vulnerable to exploitation by meiotic drive or sister killer SGEs, but to the best of our knowledge, there is currently neither theoretical nor empirical support for this idea. Another possibility is that meiosis I tends to be reductional because it allows for DSB repair by sister chromatid exchange in arrested female meiosis [66]. Alternatively, the order of meiotic divisions could merely be a ‘frozen accident’, i.e., a solution that has been arrived at a long time ago by chance, and that reversal is difficult (at least with monocentric chromosomes). However, a recent paper investigating human female meioses in unprecedented detail casts doubt on this view [67]. The careful genotyping of eggs (or embryos) and polar bodies at many markers indicated that surprisingly often, chromosomes followed an ‘inverted meiosis’ pattern of segregation, even though this led to aneuploidies in ~23% of cases. The question of why one order of meiotic divisions is almost universal therefore remains unresolved.

### 2.3 Meiosis modifications and loss of sex

Many organisms have abandoned canonical sexual reproduction, reproducing asexually by suppressing or modifying meiosis and producing diploid eggs that can develop without fertilisation. This raises two connected mysteries: why are some types of modifications much more frequent than others, and how can mitotic (or mitosis-like) asexual reproduction (“apomixis” or “clonal parthenogenesis” in animals, “mitotic apomixis” in plants) evolve from meiosis? Examples of meiosis-derived modes of asexual reproduction include chromosome doubling prior to meiosis (“endomitosis” or “pre-meiotic doubling”), fusion of two of the four products of a single meiosis (“automixis” in animals, “within-tetrad mating” in fungi), and suppression of one of the two meiotic divisions (included under “automixis” or “meiotic apomixis”, depending on the author; see [68–74] for detailed descriptions of these processes).

Two particularly common modes of asexuality are the suppression of meiosis I, and automixis involving fusing meiotic products that were separated during meiosis I (“central fusion”, Fig. 2F). Both are genetically equivalent and lead to reduced heterozygosity when there is recombination between a locus and the centromere of the chromosome on which it is located. Most other forms of meiosis-derived asexual reproduction lead to a much stronger reduction in offspring heterozygosity [75–79], and it has been hypothesised that the reduced fitness of homozygous progeny explains the rarity of these other forms [71,78,80]. Indirect support comes from the observation that species with regular asexual reproduction usually do so by central fusion or suppression of meiosis I, often accompanied by very low levels of recombination, thus maintaining heterozygosity. In contrast, species that only rarely reproduce asexually show a wider variety of asexual modes and higher levels of recombination [1,71,73,81,82]. Nonetheless, this hypothesis cannot explain some observations, for instance the rarity of pre-meiotic doubling with sister-chromosome pairing, which would also efficiently maintain heterozygosity [71]. Perhaps evolving a mechanism that ensures exclusive sister-pairing (i.e. the complete absence of non-sister pairing) is difficult, though it seems to occur in some lizard species [83]. In addition, such a system would make it difficult to repair DSBs occurring before doubling (as both sister chromatids would have the same DSBs) [71].

The question of how a mitotic asexual mutant can invade a sexual species is at the heart of the debate on the evolutionary maintenance of sex, as this is what is investigated in most theoretical models, and is the situation where the cost of sex is most evident [1]. However, unless meiosis can be entirely bypassed (e.g. as with vegetative reproduction), secondary asexuality is likely to evolve via modification of meiosis, keeping much of the cell signalling and machinery intact [65,76,80,81, see also section 3]. Indeed, detailed cytological and genetic investigations in several asexual species thought to reproduce clonally by mitotic apomixis have uncovered remnants of meiosis [73,86–88]. In *Daphnia*, meiosis I is aborted mid-way and a normal meiosis II follows. Hence, clonality in *Daphnia* is meiotically derived [86]. This should lead to loss of heterozygosity in centromere-distal regions, but if recombination is fully suppressed the genetic outcome resembles mitosis. Importantly, this suggests a possible stepwise route to evolution of mitosis-like asexuality. Rare automixis (spontaneous development of unfertilised eggs) occurs in many species [1,81]. If this becomes more common, forms of automixis maintaining heterozygosity in centromere regions might be selectively favoured and recombination suppressed, eventually leading to meiosis-derived asexuality with the same genetic consequences as mitosis [84,85,89–91]. Indeed, in *Arabidopsis*, meiosis can be transformed to genetically resemble mitosis, but modification of several genes is needed to achieve this [92–94]. In angiosperms, there is also the difficulty to overcome the absence of endosperm fertilization to achieve proper seed development, which further stresses that meiosis-derived asexuality is unlikely to evolve in a single step. To fully understand the evolutionary maintenance of sex, we may therefore need to understand the selection pressures acting in the intermediate stages, which probably involve loss of heterozygosity, and thus inbreeding depression [77,80]. In many cases, the initial evolution of asexuality may thus resemble the evolution of self-fertilisation, and several traits may pre-exist (such as low recombination rates) that make the successful transition to asexuality more likely in some taxa.

## 3. Meiosis punctuates life cycles

Meiosis is a key step in sexual life cycles, as well as some asexual life cycles derived from sexual ancestors. In multicellular eukaryotes, where meiosis is tightly associated with reproduction (unlike in many protists), meiosis is also a cellular and genetic bottleneck at the critical transition between the diploid and the haploid phases.

### 3.1 Meiosis timing and arrest

In early haploid eukaryotes, meiosis probably quickly followed endomitosis or syngamy. Today, multicellular eukaryotes exhibit a variety of life cycles in which the haploid or diploid phase may predominate. The duration of the different phases was perhaps initially controlled in part by the timing of meiosis-for instance, a multicellular, extended diploid phase likely evolved by postponing meiosis. However, in metazoans, life cycles are mostly determined by the extent of somatic development within each phase rather than by the timing of meiosis, which can be halted or postponed. In animals, where haploid mitosis is suppressed, syngamy immediately follows meiosis. Furthermore, specific cells are ‘destined’ at an early stage to eventually undergo meiosis (a.k.a. germline), whereas this cell fate is determined much later in fungi, plants and some algae.

The timing of meiosis in the germline of animals has been intensively investigated. Whereas male meiosis occurs continuously, female meiosis usually stops twice (Fig. 1). These ‘meiotic arrests’ are under the control of various factors that are not completely identified across animals [95–97]. Arrest 1 occurs in prophase I during early development and can last years until sexual maturity. The timing of arrest 2 is more variable (ranging from metaphase I in many invertebrates, to metaphase II in vertebrates and G1 phase after meiosis II in some echinoderms), and may have evolved to prevent the risk of premature parthenogenetic cleavage of oocytes or inappropriate DNA replication before fertilisation [97,98]; this is supported by the fact that this arrest is usually released by fertilisation. However, the evolutionary significance of its precise timing in diverse groups is not well understood. Three ideas have been put forward to explain arrest 1 [66]. First, its occurrence at prophase I may allow the repair of accidental DSBs by sister chromatid exchange during long periods between arrests 1 and 2. Second, if arrest 1 was to occur during an earlier mitotic division within the germline, this might decrease the variance in the number of deleterious mutations among gametes within individuals, which may be detrimental if some defective gametes or early embryos can be eliminated and replaced during reproduction. Third, it may be easier to prevent uncontrolled proliferation in a non-dividing meiotic oocyte, as once the cell starts the meiotic cell division, it cannot engage in further mitotic divisions. Arrest 1 may thus have evolved to control (and minimise) the number of possibly wasteful and mutagenic mitotic divisions in the female germline. Similar meiotic arrests in plants are unknown. Plants seem to completely lack strict mechanisms to arrest the meiotic cell division. Contrary to animals and fungi that may arrest the cell cycle and abort meiosis once DSBs are not repaired, plants will progress through meiosis irrespective of such major defects [40].

### 3.2 Meiosis and epigenetic reset

Meiosis and syngamy represent critical transitions between haploid and diploid phases in each generation. It has been suggested that a primary function of meiosis is to allow for epigenetic resetting in eukaryotes [99]. For instance, metazoan development is under the control of many epigenetic changes (cytosine methylation and chromatin marks) that are irreversibly maintained throughout life and must be reset twice each generation (at the n →2n and 2n→n transitions). This ensures proper development, the acquisition of parent-specific imprints, and may allow for mechanisms limiting the maximal number of possible successive mitoses (“Hayflick limit”, reducing tumour development [99]). Some loci escape these resets, which can lead to transgenerational epigenetic inheritance [100]. This occurs much less frequently in animals than in plants (e.g. in *Arabidopsis*, demethylation is largely restricted to asymmetric CHH methylation sites, and contrary to mouse, does not occur on most symmetric CG and CHG methylation sites) [100]. Although the 2n→n resetting occurs at or very close to meiosis in some cases (in female meiosis in animals), its timing may not be strictly tied to meiosis. For instance, it occurs pre-meiotically in the male germ line of animals (as shown in mice) or post-meiotically in male plant gametophytes (as shown in *Arabidopsis*) [100].

The evolutionary significance of these timing differences are poorly understood. Meiosis may simply not be the optimal time for epigenetic resetting. Many epigenetic pathways repress the activity of transposable elements (TEs), and so resetting epigenetic marks exposes the genome to mobilisation of these elements, which may be particularly detrimental when producing gametes. In addition, meiosis may be specifically vulnerable to TE activity for several reasons [101,102]. These include (i) deficient synapsis and repair due to the reshuffling of the meiotic machinery towards TEs-induced DSBs; (ii) ectopic recombination among TEs; and (iii) interference with synapsis due to TE transcriptional activity. Alternative TE silencing mechanisms, such as those involving small RNAs, may have evolved to ensure proper TE control during epigenetic resetting. For example, these mechanisms involve piRNA and/or endo-siRNA in mammal male and female germlines, respectively [103], and transfer of siRNA from the central cell to the egg cell in plant female gametophytes [104]. It is also possible that stringent synapsis checkpoints evolved, in part, to prevent the formation of defective gametes due to TE activity, along with other possible causes of meiotic errors.

### 3.3 Meiosis asymmetry

Symmetrical meiosis results in four viable gametes, whereas asymmetrical meiosis results in a single gamete. Symmetrical meiosis is ancestral and is found in male meiosis in animals, seed plants, ‘homosporous’ species (e.g. mosses, many ferns) and isogamous eukaryotes. Asymmetrical meiosis, on the other hand, has evolved multiple times, and occurs in female meiosis in animals, seed plants and some ciliates. The selective scenarios underlying the evolution of meiotic asymmetry are unresolved. In some cases, such as in ciliates, there is no requirement for four meiotic products, as sex occurs by the cytoplasmic exchange of haploid micronuclei (“conjugation”). In other cases, asymmetrical meiosis in females results in a large oocyte full of resources, which may favour the production of a single cell rather than four [66,105,106]. However, females could also achieve this symmetrically by undergoing fewer meioses. Therefore, is is possible that asymmetrical meiosis allows better control of resource allocation to oocytes, as symmetrical meiosis may not ensure an even distribution of resources across four meiocytes; one difficulty here is that it is not clear why female control of resource allocation would be more efficient among meiocytes derived from the same or different meiosis. A solution may be that meiocytes must compete for resources during meiosis, so that a symmetrical female meiosis is vulnerable to SGEs that bias resource allocation in their favour, possibly by killing other products of meiosis [106]. Asymmetrical meiosis may therefore have evolved to suppress such costly competition within tetrads [107], but as discussed in the next section, it also opens the possibility of new conflicts [106]. Hence, the evolution of asymmetrical female meiosis is a question that remains not entirely resolved.

### 3.4 Fairness of meiosis

A striking feature of meiosis is its apparent fairness: under Mendel's first law of inheritance, each allele has a 50% chance of ending up in any given gamete. However, there are many SGEs that increase their chances above 50% by subverting the mechanism of meiosis. These SGEs fall into two classes. The first class is killer SGEs, which kill cells that have not inherited the element. In principle, such killers could operate during meiosis (the hypothetical 'sister killers' as discussed above), but the numerous killer SGEs that have been identified so far operate postmeiotically, e.g. by killing sibling sperm [108–111]. The second class consists of meiotic drivers that exploit the asymmetry of female meiosis discussed in the previous section. These elements achieve transmission in excess of 50% by preferentially moving into the meiotic products that will eventually become the eggs or megaspores [109,112]. There is a similarity between this kind of meiotic drive, where alleles preferentially go where resources are (i.e. the egg), and SGEs expressed later and biasing resource allocation in their favour [113]. Parents make decisions of allocations to offspring before the “meiotic veil of ignorance”, whereas offspring compete for resources “from behind the veil” [114,115]. These genetic conflicts (between parent and offspring and between paternally and maternally derived alleles) are likely at the origin of parental imprints that differentially occur at male and female meiosis on some genes controlling embryo growth [114].

SGEs that undermine the fairness of meiosis provide explanations for otherwise puzzling observations. Perhaps most strikingly, centromere DNA regions often evolve rapidly, in contrast to what one would expect given their important and conserved function in meiosis. Henikoff *et al.* [116] therefore proposed that expansion of repeat sequences in centromeric DNA produces a “stronger” centromere, with increased kinetochore binding, which exhibits drive towards the future egg during meiosis I and consequently spreads in the population. Some of the best support for this hypothesis comes from a female meiotic driver in the monkeyflower *Mimulus guttatus* [117], Although conclusive evidence for a direct centromere function of this element is lacking, it is physically associated with large centromere-specific satellite DNA arrays [118]. Female meiotic drive may also explain rapid karyotype evolution and the distribution of meta-vs. acrocentric chromosomes [112] because Robertsonian fusion chromosomes (fusions of two acrocentric chromosomes into one metacentric) can behave like meiotic drivers and segregate preferentially into the future egg during meiosis I [119].

Other features of meiosis may be adaptations to suppress killer or meiotic drive SGEs. Such adaptations are expected, because these elements are generally costly for the rest of the genome [e.g. 108,120]. Defence against killer elements can be achieved by limiting gene expression. Accordingly, meiotic sex chromosome inactivation (MSCI, starting at pachytene of prophase I, see Fig. 1) has been proposed to have evolved to control sex chromosome meiotic drive elements [121], and more generally this same principle may explain limited gene expression during meiosis and in its haploid products, as well as sharing of RNA and proteins among these cells. There is also evidence for rapid evolution and positive selection in the DNA-binding regions of centromere-associated proteins, which accords with the expectation of selection for countermeasures to limit preferential segregation of centromere drive elements towards the egg [106,116]. The evolution of holokinetic chromosomes may be an extreme form of defence against centromere drive [106].

## 4. Meiosis and recombination

A ubiquitous feature of meiosis is the exchange of genetic material between homologous chromosomes. Whilst we have discussed arguments on its origin (see section 1.2), the maintenance of recombination is even more debated [122–124]. Here, we do not review this question, but discuss the evolutionary significance of patterns of recombination variation within and across species, as these present many mysteries connected to the functioning of meiosis.

### 4.1 The number of crossovers per chromosome: constrained or not?

In many species, the number of COs per bivalent appears to follow highly constrained patterns, showing little variation compared to the variation of chromosome sizes, themselves spanning several orders of magnitude [125]. Within species, the correlation between genetic map length (in cM, with 50 cM being equivalent to 1 CO per bivalent) and physical length (in megabases, Mb) per chromosome is very strong (R^2^>0.95) [126–131], and often has an intercept of ~50 cM, consistent with occurrence of one obligate CO per bivalent. There is direct evidence indicating that bivalents lacking a CO have an increased probability of non-disjunction, resulting in unviable or unfit aneuploid offspring [132,133]. Indeed, COs establish physical connections between homologues, promoting accurate disjunction by providing the tension needed for the bipolar spindle to establish [134–136]. Therefore, this constraint has likely led to the evolution of regulation of CO numbers per bivalent across the eukaryotes [137,138]. However, the reasons underlying the evolutionary persistence of this constraint are not well understood. In several species [e.g., *Arabidopsis*, 139], the intercept is less than 50cM, but the smallest chromosome is at least 50cM, thus still consistent with one obligate CO. More decisively, many species are achiasmate (i.e. have an absence of recombination) in one sex [140], with alternative mechanisms to ensure proper disjunction of achiasmate bivalents [141,142]. This indicates that COs are not always obligatory and are maintained for reasons other than ensuring proper disjunction.

In addition to the obligate CO, additional CO events can occur within bivalents. The strong cM-Mb relationship within species indicates that the number of surplus COs correlates strongly with physical chromosome size (see above). However, the rate at which surplus COs are added per Mb (i.e. the slope of the correlation) varies strongly between species [125,131,143]. This may be partly explained by selection for different CO rates in different species [144–146]. The strong correlations observed within most species may be explained by variation in trans-acting factors, such as the locus *RNF212* and its protein, which affects the propensity for DSBs to form surplus COs [147,148]; indeed, the identification of loci affecting variation in CO rates indicates the potential for rapid evolution of CO rates within and between species [149].

A further constraint on bivalent disjunction may exist: the separation of different bivalents on the meiotic spindle may need to be collectively synchronised to avoid aneuploidy. If the number of COs correlates with the amount of tension exerted on the homologues, then a tight control of excess COs may minimise disjunction asynchrony. This hypothesis may explain the observation that some disjunction problems in humans occur in a global manner without involving effects driven by specific chromosomes [150–152]. Generally high CO numbers are, on the other hand, not necessarily problematic with respect to proper disjunction [153,154].

### 4.2 Crossover interference

A CO in one position may strongly reduce the likelihood of another CO occurring in the vicinity and/or on the same bivalent. This ‘crossover interference’ is widespread [125,153,155,156], but its function and mechanistic basis remains largely unknown. In many species, two classes of COs have been identified: Class I COs, which are sensitive to interference; and Class II COs, which are not [157]. Class I COs are thought to play a major role in ensuring obligate COs, and so interference may limit the frequency or variance of COs, which may be important in ensuring proper disjunction [158]. For instance, as with autopolyploids (see above), increased interference may limit the number of CO to just one per chromosome, preventing aberrant multivalent segregation [54]. A variant of this idea is that interference is a mechanism to avoid COs occurring in close proximity, which might reduce cohesion between homologues [159] or slip and cancel each other out when they involve 2 or 4 non-interlocking chromatids, resulting in no CO occurring [160]; however, these mechanisms do not explain long-distance interference. A further suggestion is that CO interference may be adaptive by breaking up genetic associations. First, adjacent COs may be avoided because they cancel their effects on genetic associations [161]. Second, it has been speculated that CO interference may reduce the chances of breaking up coadapted gene complexes (supergenes) [162]. Some support for the idea that CO interference is not a purely mechanistic constraint comes from the fact that some species lack interference [155] and, more importantly, that there is some evidence suggesting that interference levels evolve in long-term evolution experiments in *Drosophila* [163].

### 4.3 Differences in recombination rates between the sexes

In many species, CO rates and localisation differ between male and female meioses, and these differences can vary in degree and direction even between closely related species [164–166]. The most extreme case is achiasmy, an absence of recombination in one sex, nearly always the heterogametic sex [164]. This may have evolved either as a side effect of selection to suppress recombination between the sex chromosomes [167,168], or as a way to promote tight linkage without suppressing recombination on the X or Z chromosomes [165]. More intriguing are the quantitative differences between males and females, known as heterochiasmy, which are found in many taxa, but whose mechanistic and evolutionary drivers are not yet fully understood. A number of explanations have been proposed, relating to mechanistic factors such as differences in chromatin structure [169–171], sexual dimorphism in the action of loci associated with CO rate [e.g. *RNF212*, 127,128,148], and evolutionarily widespread processes such as sperm competition, sexual dimorphism and dispersal [164,172,173]. Some models point to a role of sex differences in selection during the haploid phase [174]. Whilst a viable explanation in plants [165], there is little empirical support for this in animals [173], where meiosis in females is only completed after fertilisation (i.e. there is no true haploid phase), and where only few genes are expressed in sperm. However, meiotic drive systems are often entirely distinct between males and females [175] and may be a primary cause of haploid selection [176]. These systems often require genetic associations between two loci (a distorter and responder, or a distorter and a centromere in males and females, respectively). These driving elements might thus be very important in shaping heterochiasmy patterns [107]. Indeed, COs in female meiosis are located closer to centromeres, which would be consistent with the view that this localisation evolved to limit centromeric drive [177] (see also section 3.4). Similarly, meiotic drive in favour of recombinant chromatids have been detected in human female meiosis [67], which may limit centromere drive.

### 4.4 The localisation of COs and recombination hotspots

The localisation of recombination events differs between species. In many species, recombination occurs in localised regions known as “recombination hotspots” of around 1–2kb in length [178–181], although some species (e.g., *C. elegans* and *Drosophila*) lack well-defined hotspots [182,183]. There are at least two types of hotspots (Fig. 3). The first type, probably ancestral, is found in fungi, plants, birds and some mammals; these hotspots are temporally stable (up to millions of years) and concentrated near promoter regions and transcription start sites [180,184–187]. The second type is likely derived, and is found in other mammals, including mice and humans, where the positioning of hotspots is determined by the zinc-finger protein PRDM9. This system differs in two respects from the former: first, it appears to direct DSBs away from regulatory regions [188], and second, mutations in the DNA-binding zinc-finger array change the sequence motif targeted by the protein, leading to rapid evolution of hotspot positions over short time-scales [189,190]. This system is not present in all mammals: in dogs, hotspots target promoter regions [191], and the knock-out of Prdm9 in mouse makes recombination target promoter regions instead, underlining its derived nature [188].

**Figure 3.**
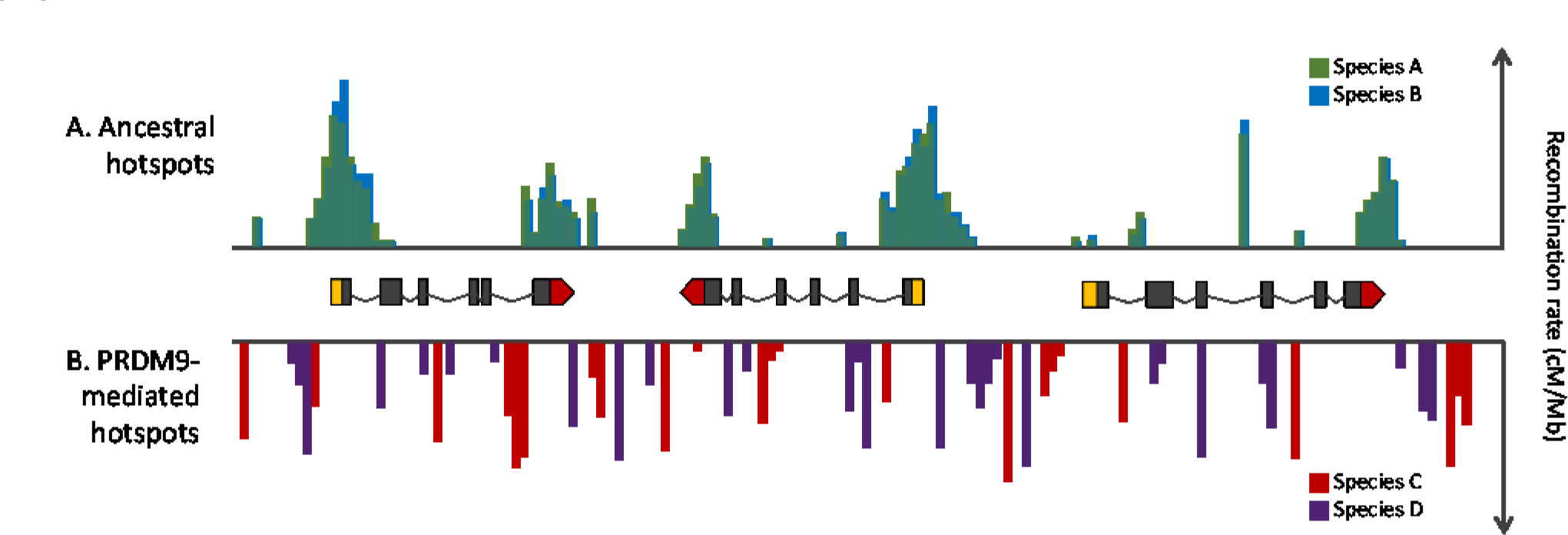
Hypothetical genome sequence containing three genes showing the distribution of ancient recombination hotspots in most model species (A) compared to derived PRDM9-mediated recombination hotspots (B). Studies in fungi, plants, birds and dogs indicate that ancestral hotspots are stable over long evolutionary timescales (up to millions of years) and concentrate at promoter regions and transcription start sites (and at stop sites in some species). These start and stop sites for each gene are indicated in yellow and red blocks, with their introns and exons represented by lines and black blocks, respectively. PRDM9-mediated hotspots are found in some mammals, including humans and mice, and are directed away from promotor regions. The DNA-binding zinc-finger in the PRDM9 protein targets specific sequence motifs; mutations in the zinc-finger array change the targeted motif, leading to rapid evolution of hotspot positions and an absence of hotspot conservation over short evolutionary time-scales (at the population and species level).

The evolutionary significance of both kinds of hotspots remains unclear. For the first type, the positions of hotspots may be caused by chromatin accessibility in transcribed regions, or, have evolved to favour recombination in gene rich regions (where it might be worth reducing genetic association). However, this does not clearly account for their precise location in regulatory regions. Another possibility might be that the co-occurrence of both COs and gene conversion events (i.e. where resolution of DSBs without CO is achieved by exchanging small segments of DNA) specifically in regulatory regions could repress enhancer runaway, a mechanism that can lead to suboptimal expression levels [192]. The evolutionary significance of the second kind of hotspot is similarly elusive. These hotspots are self-destructing because the target sequence motifs are eroded by biased gene conversion (BGC) during DSB repair [193]. This leads to a “hotspot paradox”: how can hotspots and recombination be maintained in the long term in the face of BGC [194]? A possible solution is that trans-acting factors like PRDM9 may mutate sufficiently fast to constantly ‘chase’ new and frequent targets (hotspots), switching to new ones when these targets become rare due to BGC [195]. This ‘Red Queen’ model does not require strong stabilising selection on the number of COs, and closely mimics the pattern of hotspot turnover observed in some cases [196]. However, this model does not explain how the second kind of hotspots evolved in the first place, as when it arose proper segregation was presumably already ensured by the first kind of hotspots (which, as seen in mice, are still active). Also, it does not explain why PRDM9 action is self-destructing: there is no necessity to induce DSBs exactly at the position of the target sequence for a trans-acting factor. In fact, there is no logical necessity to rely on a target sequence to maintain one CO per chromosome, as fixed chromosomal features could serve this purpose. It is worth noting here that recruiting promoter sequences for this purpose (as found for hotspots of the first kind) would be very efficient, as these sequences are highly stable and dispersed in the genome on all chromosomes. There is also no evidence so far that targeted binding motifs of PRDM9 correspond to some selfish genetic elements whose elimination would be beneficial. Overall, while spectacular progress has been made recently in elucidating hotspot mechanisms in detail (and patterns in recombination landscapes), there are still major gaps in our understanding of their evolutionary significance.

## 5 Conclusions

The evolutionary significance of meiosis has often been interpreted in an oversimplified manner, restricted mainly to the direct (DSB repair, proper disjunction) or indirect (genetic associations) effects of meiotic recombination. Yet, many features of meiosis are unlikely to be explained by effects of recombination alone, and the fields of cellular and molecular biology are uncovering new meiotic features at high rate. One of the main take-home messages of this review is that many, if not most features of meiosis are still awaiting an evolutionary explanation. Nonetheless, the recent advances in all detailed aspects of meiosis now offer the chance to investigate these questions in a far more comprehensive manner. This will require continued dialogue between cell, molecular, and evolutionary biologists [as advocated e.g. in 197], and perhaps also the realisation that similarities between features may in fact have different evolutionary explanations (e.g., different kinds of hotspots).

One of the most salient themes in most meiosis mysteries is the impact of genetic conflicts and SGEs. As for the evolution of genome size and structure, their impact is probably central [198], but in many cases, they remain hypothetical and difficult to demonstrate and study directly: many SGEs reach fixation quickly and leave almost no visible footprint. Showing that some meiotic features evolved to control SGEs represents an even greater challenge. Indeed, if successful, such features would prevent these SGEs to spread, further limiting their detection. In addition, demonstrating a role in SGEs control requires to rule out that these features evolved for more mechanistic and simpler alternatives. This is usually extremely difficult, as many *ad hoc* mechanistic constraints can be imagined.

Although meiosis is highly conserved in eukaryotes, deviations from the norm are ubiquitous and may provide important insights into its evolution. This is already apparent when considering model organisms (e.g., point centromeres in yeast, achiasmy in male *Drosophila*, holokinetic chromosomes in *C. elegans*, fast evolving recombination hotspots in mice and humans). However, the true diversity of meiotic features is likely to be revealed only when considering non-model organisms, and unicellular eukaryotes appear especially promising in this respect. Obtaining a clearer understanding of the evolutionary significance of the myriad of meiotic features will certainly be crucial to inspire and guide mechanistic investigations. Conversely, as often, *“all theory is grey, but green is the tree of life”* [Goethe, Faust Part I], and the mysteries of meiosis call for new developments of evolutionary theory, to make it less grey and more closely connected to the biological details. Overall, all these mysteries tend to have been overshadowed by the famous question of the maintenance of sex. However, resolving them might provide decisive steps towards solving this major question of evolutionary biology.

## Authors’ Contributions

T.L. conceived and co-ordinated the review. All authors contributed to and approved this manuscript.

## Acknowledgement

We thank M. Archetti, K. Bomblies, D. Charlesworth, B. de Massy, L. Duret, E. Jenczewski, R. Mercier, P. van Dijk and J. Wilkins for comments on this manuscript. Our intention was to cover all meiosis-related evolutionary mysteries in a single perspective. We apologise to all authors of the many interesting and relevant papers we were unable to cite in the final condensed version.

